# The Development of a Pipeline for the Identification and Validation of Small-Molecule RelA Inhibitors for Use as Anti-biofilm Drugs

**DOI:** 10.1101/2020.07.27.224345

**Authors:** Donald C. Hall, Jarosław E. Król, John P. Cahill, Hai-Feng Ji, Garth D. Ehrlich

**Author notes:** Corresponding Authors, **Materials & Correspondence**, Correspondences should be address to Garth D. Ehrlich and Hai-Feng Ji. Materials and strain requests should be addressed to Jarosław E. Król.

## Abstract

Biofilm infections have no effective medical treatments and can only be disrupted via physical means. This means that any biofilm infection that is not addressable surgically can never be eliminated and can only be managed as a chronic disease. Therefore, there is an urgent need for the development of new classes of drugs that can target the metabolic mechanisms within biofilms which render them recalcitrant to traditional antibiotics. This antibiotic recalcitrance of bacterial biofilms can be attributed largely to the formation of persister cells within the biofilm structure. These biofilm persister cells can be resistant to up to 1000 times the minimal inhibitory concentrations of many antibiotics as compared to their planktonic envirovars; they are thought to be the prokaryotic equivalent of metazoan stem cells. Their metabolic resistance has been demonstrated to be an active process induced by the stringent response that is triggered by the ribosomally-associated enzyme RelA in response to amino acid starvation. This 84-kD pyrophosphokinase produces the “magic spot” alarmones, collectively called (p)ppGpp. These alarmones act by directly regulating transcription by binding to RNA polymerase. These transcriptional changes lead to a major shift in cellular function to both upregulate oxidative stress-combating enzymes and down regulate major cellular functions associated with growth and replication. These changes in gene expression produce the quiescent persister cells. In this work, we describe a hybrid *in silico*-laboratory pipeline for identifying and validating small-molecule inhibitors of RelA for use in the combinatorial treatment of bacterial biofilms as re-potentiators of classical antibiotics.

## Introduction

Biofilms can be defined as a multicellular stage in the bacterial life cycle wherein bacteria through multiple intercellular communication systems create densely populated communities embedded within a self-extruded extracellular polymeric matrix(1–5). Biofilms can be resistant to antibiotic concentrations that are greater than 1000-times higher than their planktonic counterparts(6–8); moreover they also display the ability to live in extreme pHs(9). These attributes make biofilm infections extremely difficult to eradicate and nearly impossible if they are not accessible to physical means of disruption(3–5, 10, 11). The activation of the ribosomally-associated RelA enzyme *via* amino acid starvation triggers the bacterial stringent response that leads to the phenotypic changes that underlie the extreme recalcitrance that biofilms exhibit towards antibiotics(12). Thus, this ancient bacterial stress response produces an active metabolic state that results in the inability to treat chronic infections resulting from biofilms.

It has been known for more than half a century that the “magic spot” alarmones, guanine tetraphosphate and guanine pentaphosphate, collectively known as (p)ppGpp, produced by RelA, play an integral role in cell signaling for the stringent response(13–17). In *Escherichia coli*, the production of (p)ppGpp is carried out by the enzyme RelA(18). RelA, a highly conserved 84-kD pyrophosphokinase protein(19), displays a well-choreographed dance with stalled ribosomes to detect amino acid starvation by means of deacetylated tRNAs(20). Upon detection of this, uncharged tRNA, RelA subsequently binds to the ribosomal complex and structurally changes to its active synthase conformation. While in this “open” conformation, RelA continually produces (p)ppGpp(21–23). During this time, the intracellular concentrations of (p)ppGpp increase dramatically. The increased concentration of (p)ppGpp modulates multiple downstream cellular signaling pathways including interacting with the RNA polymerase’s promoter binding region, thereby interfering with the cell’s ability to produce additional ribosomes(24).

Currently, there are only a very limited number of inhibitors known for RelA and (p)ppGpp that have been identified principally through traditional drug discovery methods, such as substrate analog design and high-throughput compound screening, none of which are candidates for clinical trials for the control of biofilm infections. The first of these inhibitors were analogs to ppGpp itself, such as Relacin and its analogs(25). These compounds, while mildly effective, suffer from off-target effects and low binding affinities(25–29).

The next compound discovered to reduce the intracellular concentrations of ppGpp was the cationic peptide known as IDR1018. This peptide is an analog to bactenecin(30–33) and it was reported to directly sequester and break down (p)ppGpp, thus lowering its intracellular concentration(30). It is now thought that IDR1018 does not specifically target (p)ppGpp, but simply acts as an antimicrobial agent by means of its cationic nature(34). Moreover, IDR1018 is a moderately sized peptide incapable of being an orally administered “druggable” compound.

Recently, a trend toward the use of *in silico* chemistry and molecular modeling for computer-aided drug design has gained significant momentum(35). Previously, this was impossible to do with the RelA/RSH (RelA-SpoT homolog) family of enzymes as there were no adequate high-resolution molecular structures available. However, several RelA and related enzyme structures have been characterized and published: RelA (*E. coli*)(36), RelP (*Staphylococcus aureus*)(37), RelQ (*S. aureus*)(38), Rel_seq_ (*Streptococcus equisimilis*)(39), and Rel (*Mycobacterium tuberculosis*)(40). Thus, it has become possible through alignment and homology studies to determine the active residues within the catalytic center of these enzymes and to specifically target this region to predict and understand the ligand binding events for the rational identification of inhibitors.

## Results and Discussion

Structural modeling of the *E. coli* RelA protein(36) was performed to identify the active center. We then constructed multiple single amino acids substitution mutants of RelA based on this molecular modeling to confirm the location of the enzyme active center, and to confirm the critical role that the tyrosines Y310 and Y319 play in its enzymatic activity. Using the structural information gained from the *in silico* and laboratory studies we then developed a computationally-based pipeline to identify RelA inhibitors from large databases of known compounds that provided for the screening of compounds in a relatively timely and cost-effective manner. Millions of compounds were screened in a matter of weeks and the “hit” compounds were purchased for functional studies to determine their initial efficacy in laboratory-based *in vivo* and *in vitro* assays. The compound databases used for screening were designed to only include compounds that meet the “drug-like” criteria for ligands as defined by Lipinski’s rule of 5(41). This method has been shown to be highly effective in the discovery of drugs over the last 20 years and continues to improve in accuracy as the algorithms for ligand docking improve(42). Using these *in silico* docking studies, two small-molecule compounds that were predicted to inhibit the RelA enzyme were identified. These compounds were then subjected to *in vivo* and *in vitro* (p)ppGpp quantification assays using the *E. coli* strain C and recombinant *E. coli* RelA enzyme, as well as in biofilm inhibition assays using our *E. coli* C biofilm model(43), respectively (Fig. 1).

**Figure 1.**
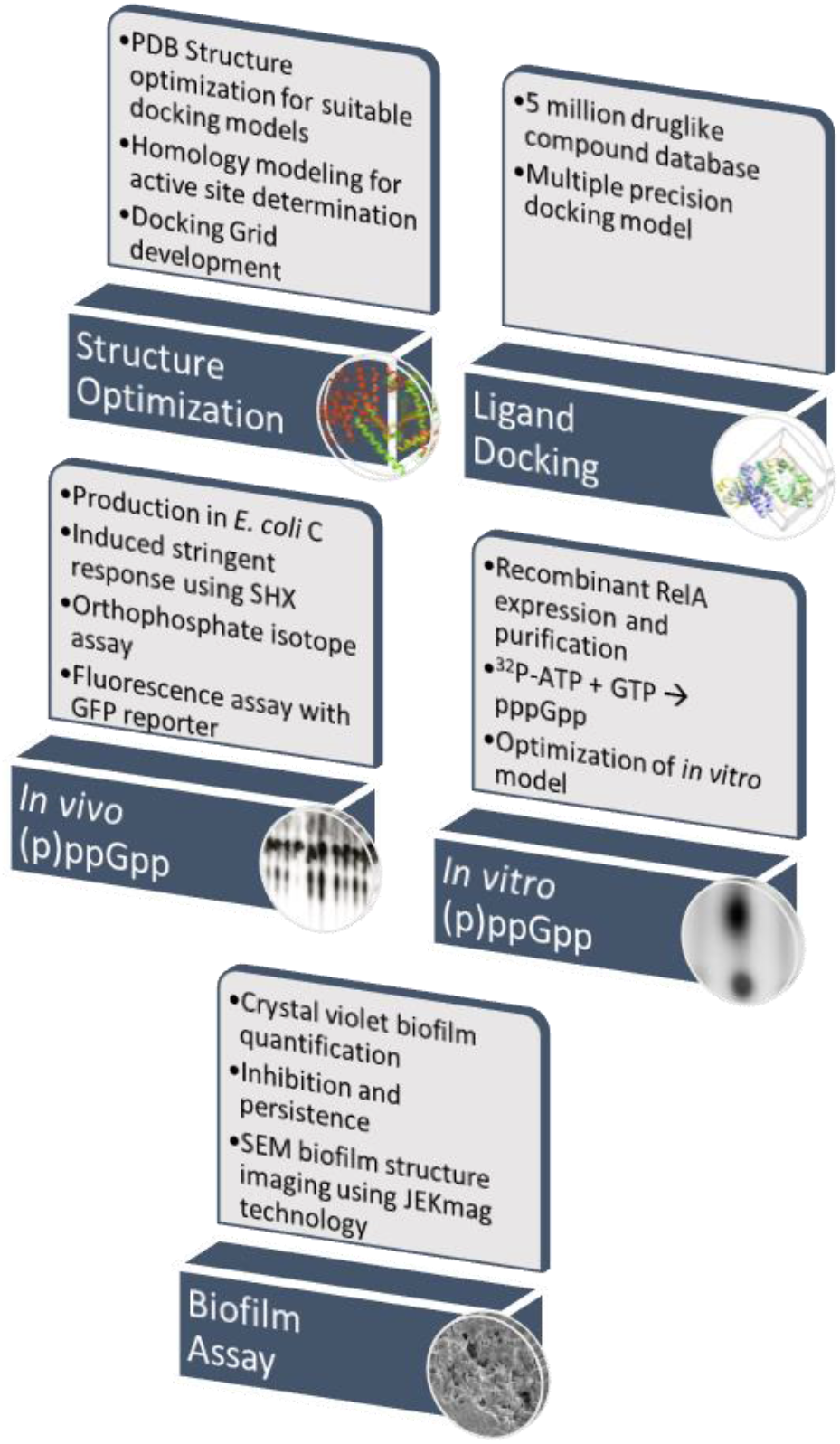
Schematic of pipeline for determination of effective RelA inhibitors.

### Validation of the RelA Activity Assays

Several methods to study the RelA enzymatic activity *in vitro* and *in vivo* have been published(25, 26, 28, 44), and our methods were adapted from these sources. We performed two kinds of RelA activity tests: a ppGpp-dependent fluorescent reporter *in vivo* assay and direct (p)ppGpp detection assays *in vivo* and *in vitro*. The first method was based on the ability of ppGpp to affect expression of different genes(45). One of these genes *rpsJ*, encodes the 30S ribosomal protein S10(46, 47). Its promoter, *PrpsJ*, belongs to the r-protein family of promoters, which are strongly inhibited by ppGpp and the DksA transcriptional factors(48, 49). Recently, a plasmid construct carrying a *yfp* (yellow fluorescent protein) gene driven by the *PrpsJ* was published(50). The reporter plasmid contains the broad host range RK2 minimal replicon and is compatible with many other plasmid vectors. Comparison of the yellow fluorescent protein (YFP) activity between wild-type (WT) *E. coli* K12 and its *relA*^−^ mutant confirmed the effect of ppGpp production on *PrpsJ* activity and served as validation of this method.

The direct (p)ppGpp detection *in vivo* and *in vitro* assays relied on different ^32^P radioactive nucleotides (γ-^32^P-ATP,α-^32^P-GTP) for use as substrates, and thin-layer chromatography (TLC) to separate the reaction products. Several methods were tested and optimized to give the best results for assessing the production of (p)ppGpp. It was found that the *in vitro* buffer system did not need to be phosphate free as previously indicated(51). It was also found that the concentration of magnesium needed to be above 5 mM for optimal synthesis of (p)ppGpp. Previous work had indicated that the 70S ribosome was needed for RelA to produce (p)ppGpp *in vitro*(52); however, we found this not to be the case. There was no difference observed with 5 mM MgCl_2_ with and without 70S (Fig. 2); therefore, it was not used in the *in vitro* reactions.

**Figure 2.**
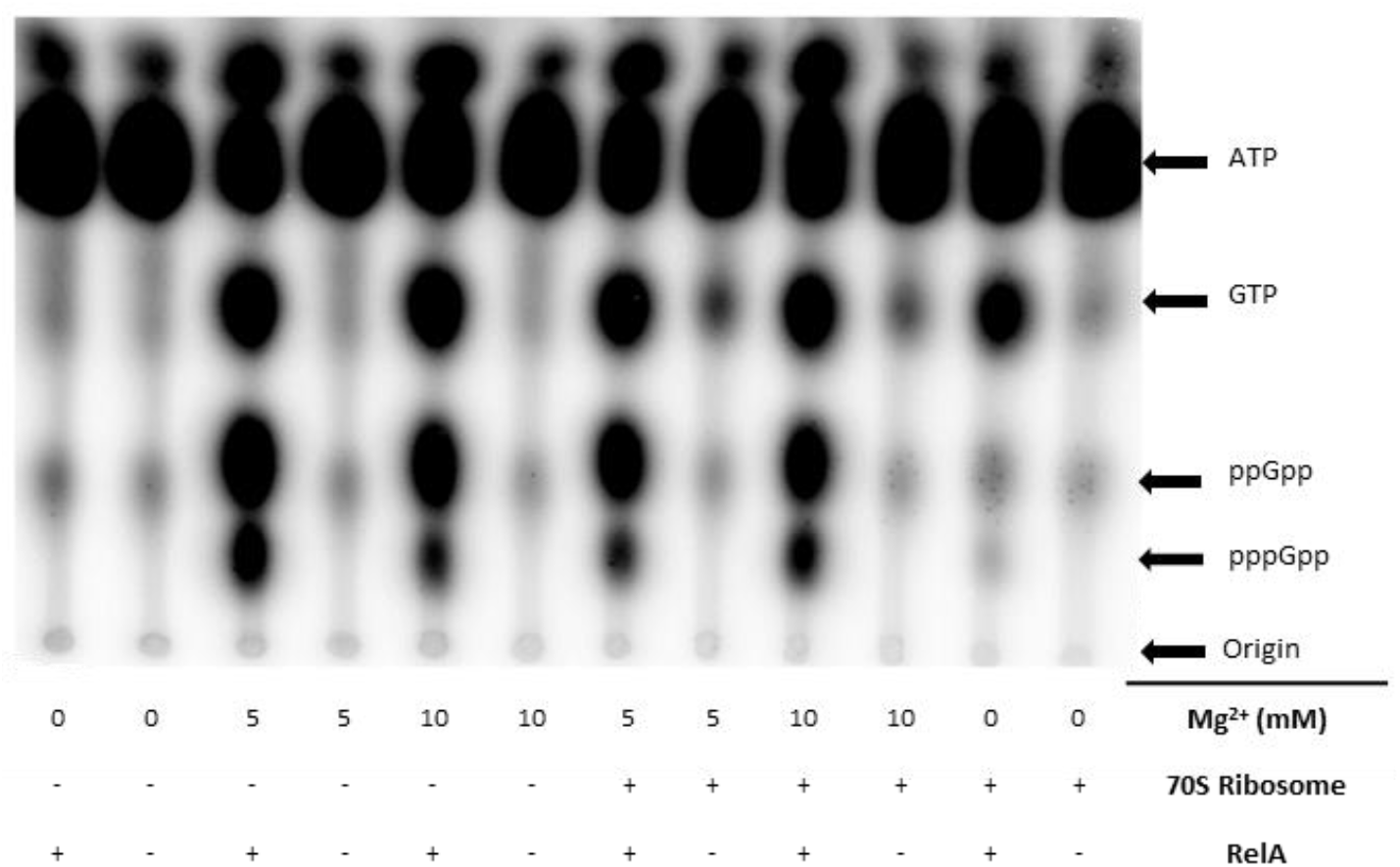
(p)ppGpp production assay. Qualitative production of (p)ppGpp in PBS buffer under various conditions: Mg^2+^ (10 mM and 5 mM), with and without 70S ribosome.

In the case of the *in vitro* assay, it was found that using γ-^32^P-ATP was optimal to study the production of both ppGpp and pppGpp, while α-^32^P-GTP was optimal for studying only pppGpp. In the case of the *in vivo* studies, [^32^P]-orthophosphate was used as the radiation source, and the cells then incorporated the ^32^P into (p)ppGpp. Both methods required TLC with a stationary phase of a polyethyleneimine (PEI)-cellulose plate and a mobile phase of 1 M potassium phosphate monobasic.

### Homology Studies

The active domain of the *E. coli* RelA cryo-EM (PDB: 5IQR) structure was determined using homology studies (Fig. S1). Because there was no substrate bound to the RelA enzyme in the cryo-EM structure(36), we utilized two methods to determine the active site for molecular docking. The first method was a genomic-based homology method, where the known RelA protein sequences were compared, and the conserved residues were evaluated (Fig. S1D).

The second method was a structural homology method in which we used crystallographic data obtained from the *S. aureus* RSH-RelP that had been co-crystallized with its nucleotide substrates to identify both the pre- and post-catalytic active sites(37). Alignment of the RelA and RelP predicted active site residues showed that they are structurally highly similar; this allowed identification and characterization of the active domain for targeting via ligand docking studies (Fig. S1A). Using this information, we were able to determine two key amino acids involved in the binding of the first substrate in the catalytic process of GDP.

### RelA Active Site Mutation Studies

To determine the accuracy of the *in silico* homology alignments and binding site determinations, two amino acid residues were identified as key to the catalytic activity of RelA, and then tested in the laboratory to ensure their assignment was correct (Fig. S1). Tyrosine residues Y-324 and Y-332 (from the alignment) (Fig. S1D) had been determined to act as one of the largest contributors to the initial binding of GDP or GTP(37). Y-324 was predicted to stabilize the phosphate of GDP/GTP by hydrogen bonding though means of its hydroxyl group; and Y-332 was predicted to be involved in π-stacking with the guanine’s aromatic ring. These stabilizations were predicted to allow for the initial binding of GDP/GTP within the active site. Y-324 and Y-332 residues correspond to the Y-310 and Y-319 residues of the *E. coli* RelA enzyme. We hypothesized that if these residues were mutated to alanines (A-310 and A-319) this should bring about a decrease in the catalytic transfer of the pyrophosphate from ATP to form (p)ppGpp. Figure S2 shows the interactions of RelA with the native residues, as well as the lack of interactions when mutated to an alanine residue. To obtain the Y/A-310 and Y/A-319 substitutions of the *E. coli* RelA, we used two synthetic DNA cassettes to replace the 5’ end of the gene in the pJW2755-AM plasmid. The first *PsiI/NsiI* (1144bp) cassette contained a silent *XbaI* mutation(53, 54)and the Y/A-310 substitution. The second 365bp *XbaI/NsiI* cassette introduced the Y/A-319 mutation (Fig. S3). The 365-bp region between the *Xba*I (769) and *Nsi*I (1144) restriction sites contain the predicted RelA active center and can be easily exchanged with a synthetic construct to replace any of the tested amino acids.

Two assays were conducted to evaluate the activity of the mutant RelA enzymes: an *in vivo* (p)ppGpp fluorescent reporter and *in vitro* (p)ppGpp production assay. The ASKA plasmid pJW2755-AM with the WT RelA protein and its Y/A-310 and Y/A319 versions were transformed into the *E. coli* AG1 strain containing a pAG001 plasmid, this plasmid contains a YFP gene expressed under a stringent response regulated promoter P*rpsJ*(50). The *E. coli* AG1 strain contains a *relA1* mutation caused by an insertion of an *IS*2 insertion sequence between 85^th^ and 86^th^ codons of the *relA* gene. These mutants retain a low level of (p)ppGpp synthesis activity(19). Plasmids pJW2755AM and its derivatives, and pAG001 belong to different incompatibility groups and therefore can co-reside in a single cell. When plasmid encoded RelA expression is induced with isopropyl β-D-1-thiogalactopyranoside (IPTG), the cells produced (p)ppGpp. Increased level of (p)ppGpp decreased the level of YPF synthesis, as it was under the control of the *PrpsJ* promoter. The results showed a much higher reduction of YFP fluorescence in the case of the WT RelA protein than with its Y/A-310 and Y/A-319 derivatives (Fig. 3A). In the *in vitro* assays the purified proteins containing the Y/A-310 and Y/A-319 when compared with the WT protein showed an even more striking reduction in pppGpp production (Fig. 3B). These results confirmed that the Y-310 and Y-319 amino acid residues play important roles in the enzymatic activity of RelA, and therefore, the active site, as modeled above, can be used as a target for the *in silico* docking of ligands for the identification of candidate, druggable inhibitors.

**Figure 3.**
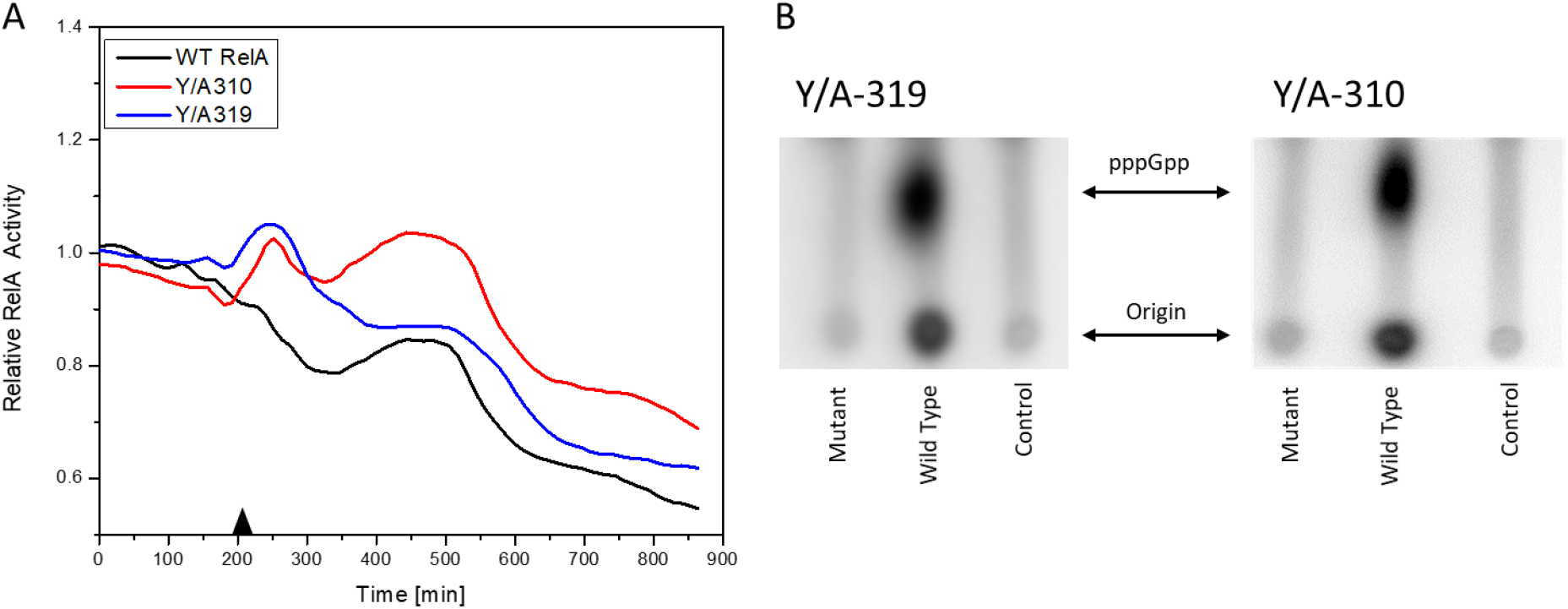
Effect of Y/A319 and Y/A310 substitutions on RelA enzymatic activity. A) Ratio of relative RelA activity of the induced to non-induced cells in an *in vivo* fluorescence assay. The induction with 1.5 μM IPTG took place at 210 min, indicated with the black triangle. B) *In vitro* pppGpp production assay. Control is [γ-32P] ATP without enzyme.

### *In silico* Screening for Hit Compounds

Non-RelA components of the *E. coli* RelA cryo-EM (PDB: 5IQR) model including RNA and ribosome were stripped away from the file leaving only the RelA structure (Fig. S4). The RelA structure was then optimized using the Schrödinger Maestro protein preparation tools including the package Prime, which allows Maestro to fill in missing side chains and determine optimal amino acid orientations. The RelA enzyme was then structurally minimized using the force field OPLS3e(55) (Fig. S4B). The enzyme binding pocket was determined using homology (Fig. S2) studies, as well as a general understanding of RelA’s function, and a docking grid box was developed for protein ligand docking calculations.

Schrödinger Maestro Molecular Modeling Glide(56, 57) was utilized to determine hit compounds, which were then validated using the laboratory assays described above to probe their ability to inhibit RelA activity. Schrödinger Glide-HTVS mode was first used to screen the entire University of California, San Francisco Zinc^12^ Database of commercially available compounds. This database contains over 4 million compounds. The top 10% from the HTVS docking scan was then filtered into Glide-SP mode (standard precision). This output was then further refined and run in Glide-XP(57) mode (extra precision). These molecular docking studies resulted in 2 compounds showing a binding score that passed our threshold for binding affinity (Table 1) and were higher than those of the natural substrates ATP and GTP. These two compounds also fit both the Lipinski’s rule of 5(58–60) for orally administered drugs, and the quantitative estimate of drug likeness(61, 62).

**Table 1.**
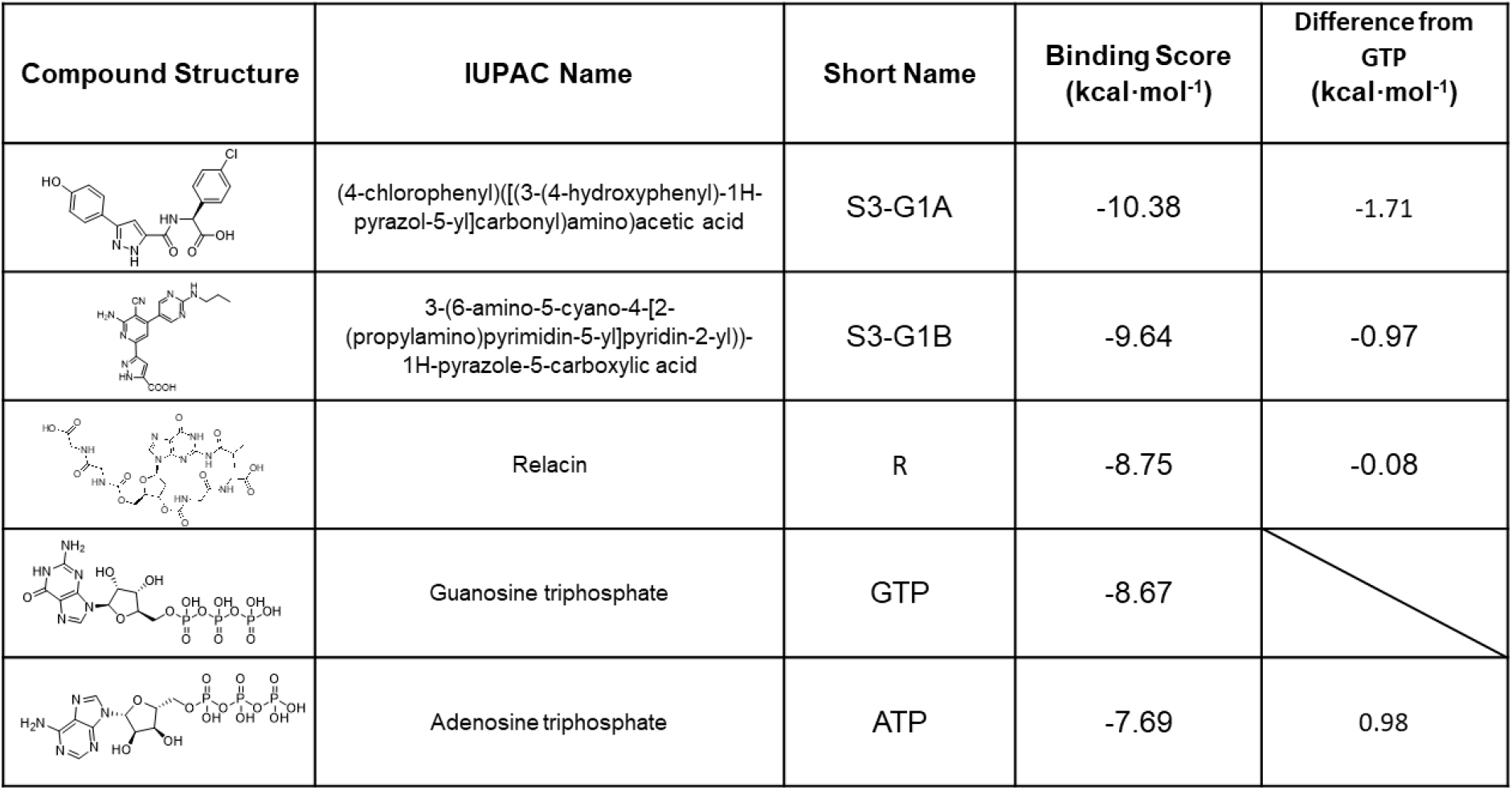
Hit compounds for the inhibition of RelA binding score. Binding score compared to the initial binding compound GTP.

### Effect of S3-G1A and S3-G1B on (p)ppGpp Production *via In Vitro* and *In Vivo* RelA Assays

After computational hit compounds were determined, the next step was to evaluate the effect of these small molecules on RelA activity in the *in vitro* and *in vivo* assays established above for the production of ppGpp. The results of the *in vitro* assay showed that both compounds S3-G1A (20 μM) and S3-G1B (20 μM) reduced the ppGpp production when compared to an untreated sample by 71.7% (*p*< 0.0001) and 79.7% (*p*< 0001), respectively (Fig. 4A). Both compounds showed higher reduction of activity than Relacin (20 μM) (45.4%, *p* = 0.0084). The *in vivo* assay showed a reduction in ppGpp production in samples treated with both compounds 31.4% (*p* = 0.0006) in S3-G1A and 17.75% (*p* = 0.0295) in S3-G1B. In this assay, no effect of Relacin on ppGpp production was observed (Fig 4B). We hypothesize that Relacin is not cell permeable and therefore does not influence the *in vivo* ppGpp production. These results indicated that the S3-G1A and S3-G1B compounds are more efficient *in vivo* and *in vitro* than Relacin and validated the entire hybrid *in silico*-laboratory pipeline.

**Figure 4.**
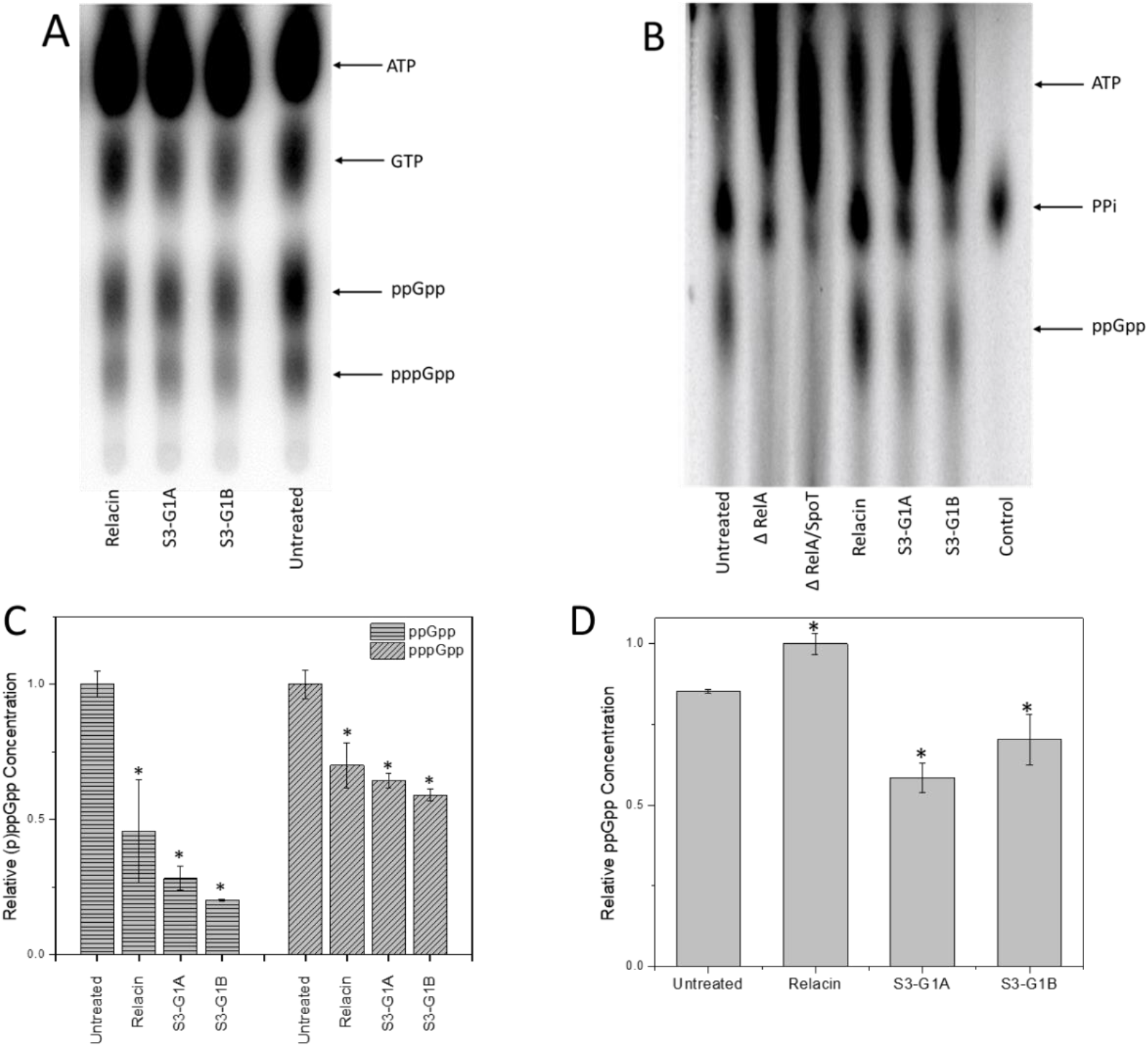
Effect of S3-G1A and S3-G1B compounds on RelA enzymatic activity. RelA (p)ppGpp production assay *in vitro* (A) and *in vivo* (B) treated with 20 μM of respective compound. Quantitative analyses of the *in vitro* (C) and *in vivo* (D) assays. * indicates statistical significance.

### Effect of Hit Compounds on Bacterial Growth

Bacterial growth rate under conditions unrestricted by substrate availability is an indicator of cell health and viability. Despite great efforts to determine the role of the stringent response on control of cell growth rate, general conclusions have not been able to be drawn(63–65). However, all reports have shown that mutants unable to produce ppGpp grow slightly more slowly (up to 30%) than their cognate WT on all media tested(63–65). We found that the initial growth rates for the WT strain and CF1652 (*relA*::Km) were the same (Fig. S5). However, growth of the WT strain started to slow down first after reaching OD_600_=0.6. The WT strain was expected to sense small changes in nutrient concentrations and react to it, reducing the growth rate. The *relA* mutant reached a higher cell density than that of the WT. After 18h of growth, both strains reached their highest cell densities and thereafter we observed varying decreases in OD_600_ values. We found that compounds S3-G1A and S3-G1B had no effect on planktonic growth rate. The maximal cell densities of the cultures with compounds were slightly lower than the control (Fig. S6).

### Effect of Hit Compounds on Biofilm Inhibition and Dispersal

We have previously reported that *E. coli* strain C(43) is the only one of the five major “laboratory strains” of *E. coli* that is a superior biofilm former; therefore, this strain was used in our biofilm assays. Studies were conducted in 96-well high-throughput assays. In the biofilm inhibition assay, compounds were added to the wells at the beginning of the experiment. For the biofilm dispersal assay, the biofilm was allowed to grow for 24 and 48h, the wells were washed with sterile phosphate-buffered saline (PBS), and fresh medium supplemented with the compounds was added to the wells. The amount of biofilm was measured after 24h. There was no observed effect on the inhibition (Fig. S7) or dispersal (data not shown) of biofilms with compounds alone.

### Effect of Compound on Biofilm Persistence and Biofilm Viability

Biofilm persistence and viability were assessed with the hit compounds in combination with an antibiotic. It has been determined that sub-MICs of antibiotic result in increased biofilm formation(66). Ampicillin was used in all of our assays due to its bactericidal effect. Sub-MIC concentrations of ampicillin were determined by growth measurements (OD_600_). We found that the biggest change in the culture cell density was observed between 40 and 60 μg·mL^−1^ ampicillin (Fig. S8A). Analyzing the effect of ampicillin on biofilm formation, we observed that the presence of the antibiotic significantly increased the amount of biofilm with the highest biomass observed at relatively high ampicillin concentrations (80 μg·mL^−1^) (Fig. S8B). To analyze the effect of our hit compounds in combination with antibiotics, a range of ampicillin concentrations from 30 to 50 μg·mL^−1^ was utilized.

The amount of biofilm biomass was determined in the combined presence of antibiotics and either compound A & B. This combination therapy led to a highly significant reduction in biofilm mass compared to the ampicillin-only-treated control (Fig. 5A). As a reference control, we used IDR 1018, an antimicrobial peptide that was reported to target (p)ppGpp directly and degrade ppGpp *in vitro*(30). Addition of the hit compounds to ampicillin concentrations of 40 μg·mL^−1^ (Amp40) and 50 μg·mL^−1^ (Amp50) resulted in a highly significant decreases in biofilm volume compared with their cognate antibiotic only control (Fig. 5A). At Amp40 the biofilm biomass was reduced by 97.9% (*p* = 0.0009) for S3-G1A (50 μM), by 92.4% (*p* = 0.0014) for S3-G1B (50 μM), and by 75.4% (*p* = 0.006) for IDR1018 (6 μM). Amp50 showed reductions in biofilm biomass of 67.9% (*p* = 0.0044) for S3-G1A, of 72.9% (*p* = 0.0042) for S3-G1B, and 65.2% (*p* = 0.0054) for IDR1018The difference between Amp40 and Amp50 can be attributed to the larger increase in biofilm volume from the higher concentration of antibiotic.

**Figure 5.**
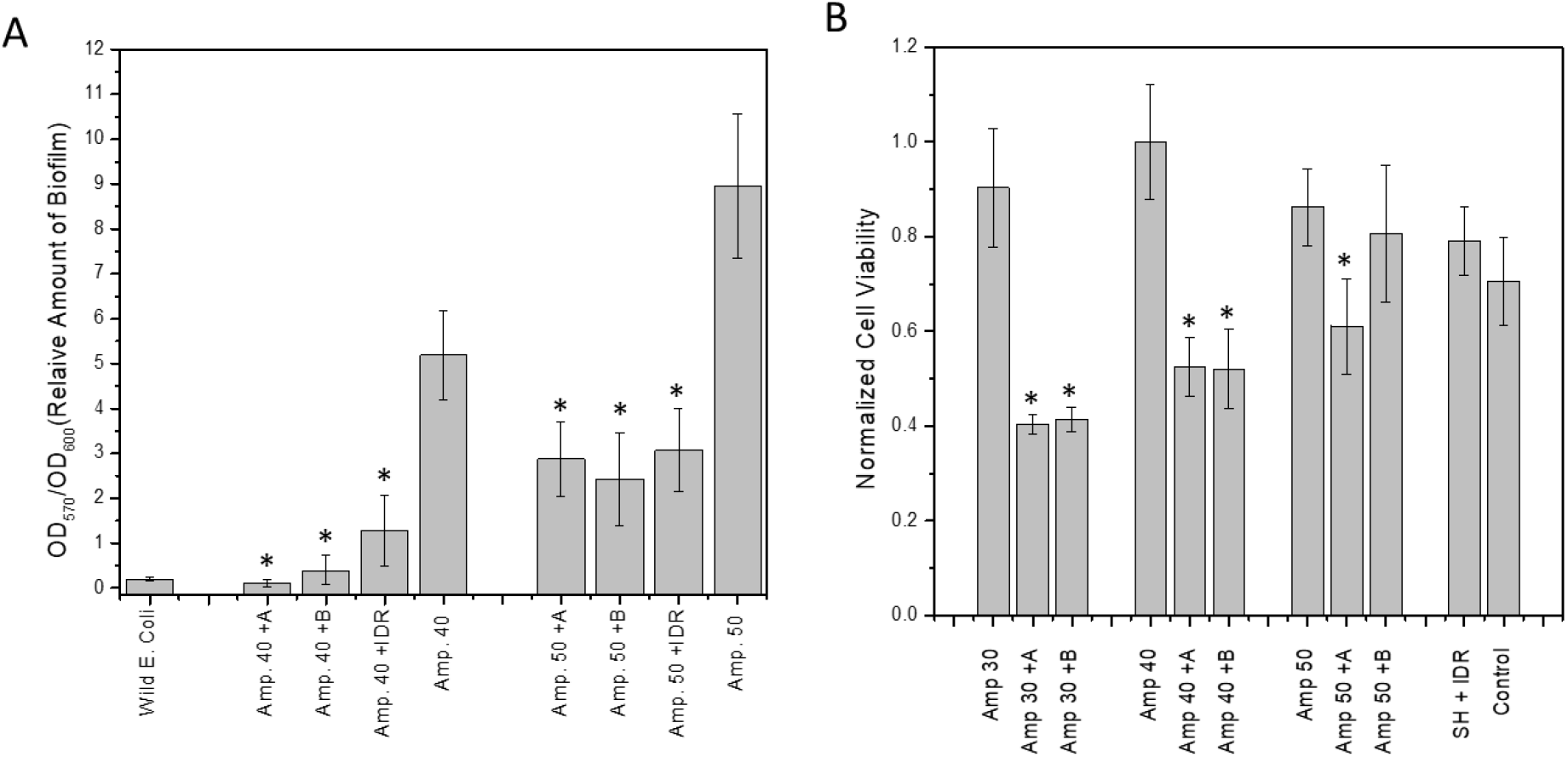
Synergistic effect of compounds and ampicillin on amount of biofilm and cell viability in biofilm. A) Biofilm degradation utilizing hit compounds and ampicillin. A = S1-G1A (50 μM), B = S1-G1B (50 μM), Amp# = Ampicillin μg·mL^−1^ SH = serine hydroxamate IDR = IDR 1018; B) AlamarBlue viability assay following combined treatment of cells with hit compounds and ampicillin shows a reduction in bacterial viability. * indicates statistical significance.

An AlamarBlue cell viability assay also showed that ampicillin killed more bacterial cells in combination with the tested hit compounds (Fig. 5B). In the case of ampicillin at 30 μg·mL^−1^ (Amp30), the reduction was 55.4% (*p* = 0.0024) and 54.2% (*p* = 0.0027) for S3-G1A (50 μM) and S3-61B (50 μM), respectively. When higher concentrations of antibiotic were used, the synergetic effects of compounds S3-G1A and S3-G1B were less noticeable, with the decreases being only 29.2% (*p* = 0.0278) and 6.5% (*p* = 0.6), respectively. This effect we attributed to the greater volume of the biofilm contained in these samples (Fig. 5B).

### Effect of Hit Compounds on Biofilm Structure

Scanning electron microscopy (SEM) allowed us to probe the structure of the biofilms treated with the hit compounds. Biofilms were grown on metal pins for 3 days that were transferred daily to fresh LB medium using the JEKMag technique(67). We found that while there was not a large reduction in biofilm mass by the compounds alone, there was a very substantial change to the structure of the extracellular matrix of the biofilms. Biofilms treated with compounds 40 μg·mL^−1^ S3-G1A and 40 μg·mL^−1^ S3-G1B exhibited a greatly reduced amount of matrix compared to untreated WT *E. coli* C (Fig. 6). Treatment with S3-G1B also resulted in elongation of the cells, indicating the possibility of an unknown off-target effect inducing filamentation.

**Figure 6.**
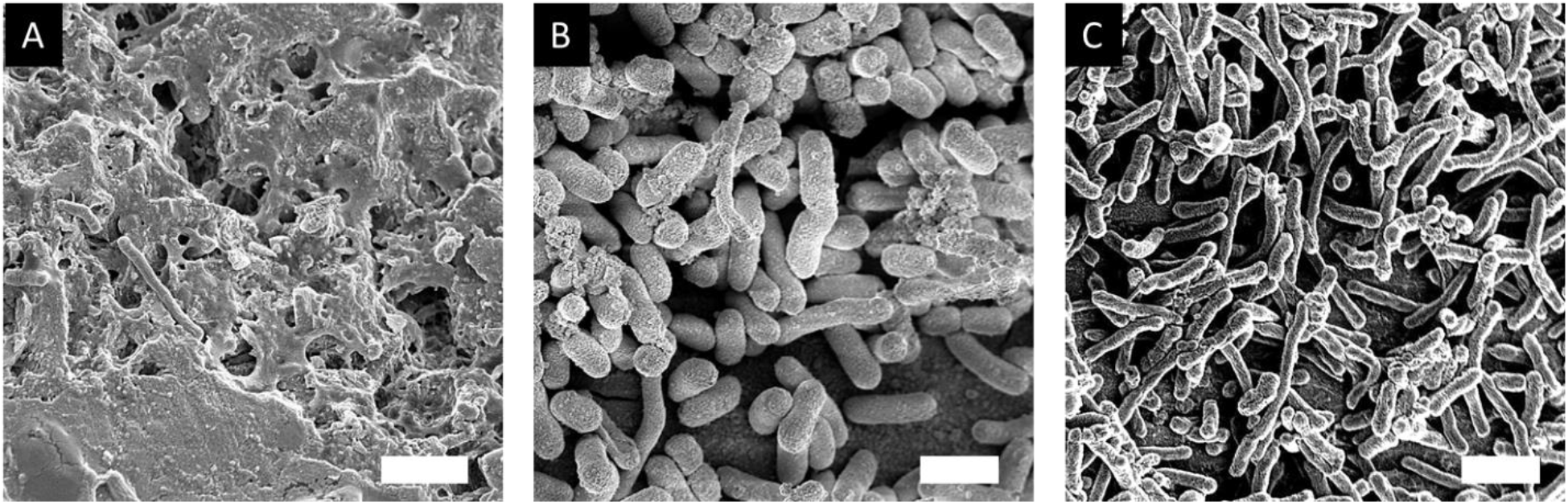
SEM images of *E. coli* C biofilm. A) Untreated wild type *E. coli* C biofilm; B) *E. coli* C biofilm treated with 40 μg·mL^−1^S3-G1A; C) *E. coli* C biofilm treated with 40 μg·mL^−1^S3-G1B. Scale bar = 1 μm

## Conclusion

We have established a hybrid *in silico-*laboratory pipeline method to identify and characterize novel RelA inhibitors for the treatment of medically relevant bacterial biofilms in combination with traditional antibiotics. Using these reported methods in combination has given us the ability to determine and characterize hit compounds from a large database of *in silico* ligands. These methods have provided us with two lead compounds that are being utilized in downstream optimization structure-activity relationships to improve the efficacy of the core bioisostere. The methods outlined here are important steps toward the process of finding an effective inhibitor of the RelA-driven bacterial stringent response and, in turn, the treatment of persistent biofilm infections. The computational components, which include binding site determinations and a multi-step docking process that incorporates a series of every more stringent filters provides for the efficient screening of large ligand libraries, and provides an effective and cost-effective means for identifying hit molecules for the inhibition of RelA. Before the addition of these *in silico* methods, high-throughput ligand assays in the biofilm space have been costly and time consuming.

## Methods

### Bacterial Strains and Growth Conditions

Bacterial strains are listed in Table S1. All bacterial strains were grown in Luria-Bertani broth (LB) or LB agar (1.5%). Antibiotics kanamycin (50 μg·mL^−1^), ampicillin (100 μg·mL^−1^), and chloramphenicol (25 μg·mL^−1^) were used when necessary. S3-G1A(68) and S3-G1B(68) were purchased from Hit2Lead and used at the concentrations described in text.

### Computational Docking

#### High-throughput *in Silico* Docking Studies

The RelA enzyme (PDB: 5IQR) was prepared and optimized using Maestro Protein Preparation (Schrödinger Maestro, New York, NY, USA; Version 11.9.011, MMshare Version 4.5.011, Release 2019-1, Platform Windows-x64). The 5IQR PDB file contained extraneous portions of the ribosome, as the structure was determined as a RelA dimer with the ribosome. The ribosome and RNA subunits were removed and RelA was isolated in a separate file. The dockable RelA structure was prepared and minimized using Schrödinger’s protein preparation application(69). This application was utilized to add hydrogens, create missing disulfide bonds, and determine lowest-energy residue orientations. Geometry minimization was carried out using the force field OPLS3e(55). A docking site was determined using homology studies of bacterial *rel* genes from several species in combination with the Schrödinger binding site determination tool. Ligands were prepared using Schrödinger LigPrep (Schrödinger Release 2020-1: LigPrep, Schrödinger, LLC, New York, NY, USA, 2020).

### Biological Validation Assays

#### RelA Mutagenesis

The ASKA(-) clone JW2755-AM containing an *E. coli* W3110 RelA in the pCA24N vector was used for mutagenesis. A 1144-bp *PsiI/NsiI* fragment was replaced by a synthetic construct. This construct contained 2 designated changes. First, a single nucleotide silent substitution (769 C/A) introduced an *XbaI* restriction site as described(53, 54). Second, a TA/GC substitution at position 1027 replaced TAC (Y-310) with GCC (A-310). The 365-bp region between *Xba*I (769) and *Nsi*I (1144) contains the RelA active center and can be easily swapped with a synthetic construct to replace any of the tested amino acids. This method was applied to introduce the Y/A-319 mutation. A 365-bp *XbaI/NsiI* fragment was replaced by a synthetic fragment with TAT-Y319 (position 1053) replaced with GCC-A319 codon as described(70). All mutations were confirmed by Sanger DNA sequencing.

#### RelA Protein Purification

the functional RelA enzyme and its Y/A-319 and Y/A-310 mutants were purified from host cell AG1 strains carrying the pJW2755-AM, pJEK2020-43, and pJEK2020-20 plasmids, respectively. One liter of LB broth was inoculated with 20 mL of overnight culture (OD_600_ = 0.9) and grown for 4 h (OD_600_= 0.8) before induction with 1.5 mM IPTG for 4 h. Cultures were spun down, washed with phosphate-buffered saline (PBS), and resuspended in lysis buffer (50 mM NaH_2_PO_4_, 300 mM NaCl, 10 mM imidazole, pH 8.0) for lysing. To that resuspension, 1 μL·mL^−1^ThermoFisher Halt™ Protease Inhibitor Cocktail (100X) was added without EDTA and cells were lysed with sonication on ice (cycles 10 s on 10 s off for a total of 3 min of sonication, 2X). Lysates were spun down to remove cellular debris. Millipore Sigma PureProteome™ Nickel Magnetic Beads were used according to modified manufacturer’s instructions. Supernatant was placed in 200 μL of nickel affinity beads for a period of 30 mins. Beads were captured on a magnetic rack and the supernatant was removed. Beads were then washed 4X with wash buffer (50 mM NaH_2_PO_4_, 300 mM NaCl, 10 mM imidazole, pH 8.0). RelA was eluted twice using 300 mM imidazole elution buffer (50 mM NaH_2_PO_4_, 300 mM NaCl, 300 mM imidazole, pH 8.0) and a final elution using 500 mM imidazole elution buffer (50 mM NaH_2_PO_4_, 300 mM NaCl, 500 mM imidazole, pH 8.0). An SDS-page gel was run to confirm presence and purity of RelA. Imidazole buffer was exchanged for PBS buffer and RelA was concentrated using Amicon^®^ Ultra-4 Centrifugal Filter Unit 30 KDa nominal molecular weight limit. Nanodrop showed an average concentration of 1 mg·mL^−1^ with a 260:280 ratio ~ 0.73.

#### Fluorescent Reporter RelA Activity Assay

The plasmid pAG001 (ampicillin 100 μg·mL^−1^) carrying a *yfp* fluorescent protein gene driven by the *PrpsJ* was used to detect the intracellular ppGpp concentrations as published(50). To validate the assay this reporter plasmid, which is based on the broad host range RK2 minimal replicon, it was introduced into *E. coli* K12 CF1648, and its *relA* mutants: (CF1652)(19), and AG1 (*relA1*) (NBRP Japan). To analyze the effect of overexpression of RelA and the Y/A310, and Y/A319 substitutions, ASKA plasmid pJW2755-AM(71) (chloramphenicol 25 μg·mL^−1^) and its derivatives pJEK2020-20 with the Y/A310 mutation and pJEK2020-43 with the Y/A319 mutation were extracted using the ThermoFisher Plasmid Mini DNA Extraction Kit, and transformed into AG1pAG001 strain (ampicillin 100 μg·mL^−1^, chloramphenicol 25 μg·mL^−1^). For the fluorescent RelA activity assay, overnight cultures of the selected strains were diluted 1:100 in fresh LB medium and 200 μL aliquots were placed into 96-well plates (Costar). The plates were placed in a Tecan Infinite M200 Pro Microplate Reader with a programmed growth cycle (18 h, 37 °C, orbital rotation 3.5). Cell density was measured at OD_600_ and YFP fluorescence activity was detected with 505 nm/535 nm (excitation/emission). Enzymatic activity was measured as Relative Fluorescence Units (RFU -YFP/OD_600_).

#### *In vitro (p)*ppGpp quantification

*In vitro* (p)ppGpp quantification was carried out using techniques similar to those previously reported in the literature(25, 26, 28, 44). RelA enzyme was purified as described above. Roughly 0.4 μg of RelA protein was added to a 1.5-mL microcentrifuge tube containing a reaction mix composed of 1X PBS, 5 mM MgCl_2_, 0.5 mM ATP, 0.5 mM GTP, 0.5 mM GDP, and 20 μCi[γ-32P]ATP (3,000 Ci mmol-1; PerkinElmer) and varying concentrations of the compound of interest. These reactions were incubated at 37°C for 1 h. The reactions were stopped by addition of 5 μL formic acid (88%). The reaction mixtures were then spotted on a stationary-phase polyethyleneimine (PEI)-cellulose TLC plate using potassium phosphate monobasic (1.5 M) as the mobile-phase. The plates were then dried, and the radiation levels were read using a Molecular Dynamics Storage Phosphor Screen. A Molecular Dynamics Storm 840 Phosphor imager Scanner was used to read the phosphor screen and ImageJ was used to process the images.

#### *In vivo (p)*ppGpp Quantification

*In vivo* (p)ppGpp quantification was carried out using techniques similar to those previously reported in the literature(25, 26, 28, 44). One milliliter of overnight cell culture of *E. coli* C was placed in 1.5-mL microcentrifuge tubes and pelleted. To this pellet was added 50 μL of a reaction mixture containing 20 μCi orthophosphoric acid and 40 μM serine hydroxamate in 1X MOPS minimal medium. The cell pellet was resuspended by gentle vortexing and placed in an incubator for 1 h. Cell growth arrest and cell lysis were completed by addition of 15 μL formic acid (88%). The lysate was then centrifuged to remove any insoluble components and the supernatant was spotted on a stationary-phase PEI-cellulose TLC plate. Plates were processed and analyzed as described above.

#### Biofilm Dispersal Assays

For biofilm formation on polystyrene surfaces, flat-bottom 96-well microtiter plates (Corning Inc.) were used. Two hundred microliters of bacterial culture (100X diluted overnight culture; approximately 10^7^ cells) in fresh LB medium was added to each well. These were allowed to grow for 24 h. The planktonic cells and medium were then aspirated, and the plates were washed twice with 1X PBS. Fresh LB with hit compounds were added to the biofilm wells. These cultures were then allowed to incubate at 37 °C overnight. Cell density were measured (OD_600_) using a Multiscan Go plate reader (ThermoFisher), and 30 μL Gram crystal violet (CV) (Remel; 3 g crystal violet, 50 mL isopropanol, 50 mL ethanol, 900 mL purified water) was applied for staining for 1 h. Plates were washed with water and air dried, and CV was solubilized with an ethanol:acetone (4:1) solution. The OD_570_ was determined from this solution, and the biofilm volume was calculated as the ratio of OD570 to OD_600_(43, 72).

#### Biofilm Inhibition Assays

For biofilm formation on polystyrene surfaces, flat-bottom 96-well microtiter plates (Corning Inc.) were used. The effect of different compounds on biofilm formation was tested by adding compounds at different concentrations to the bacterial culture (100X diluted overnight culture; approximately 10^7^ cells) in fresh LB medium. Two hundred microliter aliquots were pipetted into 96-well plates and placed for 24 or 48 h into a 37 °C incubator. The biofilm mass was measured by the CV staining assay described above.

#### Biofilm Persistence Assays with Ampicillin

Biofilms were grown for 24 or 48 h as described above. Planktonic cells were removed, and the biofilms were washed twice with 250 μL sterile PBS solution. Two hundred microliters of fresh LB medium with various concentrations of ampicillin were dispensed into the wells. After 18h of incubation at 37 °C, the volume of biofilm was measured by CV staining as described above.

#### Synergistic Effects of Compounds and Antibiotics

Biofilms were grown for 24, 48, or 72 h as described above. Planktonic cells were then removed, and biofilms were washed twice with 250 μL sterile PBS solution. Two hundred microliter aliquots of fresh LB medium with multiple concentrations of the tested compounds and ampicillin were dispensed into the wells. After 18h of incubation at 37 °C, the biofilm mass was measured as described above. For the AlamarBlue viability test, 4 μL of AlamarBlue (Invitrogen) was added and plates were incubated in a Biotek HT plate reader at 37 °C for 4 h. Cell viability was measured as fluorescence at 530/590nm (excitation/emission) versus compound concentration or initial cell density

#### Cell Growth Curves

The effect of the hit compounds on bacterial growth was tested by adding compounds at multiple concentrations to the bacterial culture (100X diluted overnight culture; approximately 10^7^ cells) in fresh LB medium. Two hundred microliters aliquots were pipetted into 96-well plates and placed into a Biotek HT or Tecan Infinite M200 Pro plate reader for 18h at 37 °C. Plates were shaken during incubation and the optical density (OD_630_ or OD_600_) was measured every 15 min.

#### Antibiotic Susceptibility Assays

For liquid cultures, the minimal inhibitory concentrations (MICs) of the antimicrobial drugs were determined using 96-well plates and the broth dilution method. Suspensions were then incubated at 37°C for 18 h in a Biotek HT plate reader (see bacterial growth). Biofilm destruction experiments were performed with different antibiotic concentrations, and cell densities were measured after 18h. Bacterial concentrations were calculated via optical density (OD_630_), and the lowest concentration causing 80% growth inhibition relative to the growth of the control was deemed to be the MIC.

#### Scanning Electron Microscopy (SEM) of Biofilm

*E. coli* biofilms were grown in LB with multiple concentrations of the hit compounds on metal pins(67). These metal pins were then washed twice in 1X PBS. The biofilm-containing metal pins were then placed in a 5% glutaraldehyde solution for 1 h. Metal pins were then dried using a gradient of ethanol from 50% to 100%, 5 min in each solution. The pins were sputter coated with gold at a thickness of 60 Å. SEM images were taken on a Zeiss Supra 50VP Scanning Electron Microscope 5 kV beam acceleration.

#### Statistical Analysis

Statistical analyses were performed using OriginPro 8.5. Relevant statistical data is included in results and discussion for each experiment. Error bars indicated standard deviation from the mean. Asterisks represent statistical significance of at least *p* < 0.05.

## Acknowledgements

This work was supported by the Drexel University College of Arts and Sciences, The Center for Advanced Microbial Processing, Drexel University College of Medicine start-up funding provided to G.D.E., NIH NIDCD R01 DC 02148, The Bill and Marian Cook Foundation, and The Coulter-Drexel Translational Research Partnership.

Thanks to Dr. M. Cashel for providing RelA and RelA/SpoT mutant bacterial strains. Thanks to Dr. A. Gawin for plasmids pAG0001. We thank Jocelyn Hammond and Jerica Wilson for proofreading and comments.

## Author contributions

D.C.H.J. and J.E.K. designed and performed the experiments and the analyses and wrote the original draft and edited manuscript. J.P.C. performed molecular docking studies and analyses. G.D.E. and H-F.J. conceived and supported the project and edited the manuscript. All authors read and approved the final version of the manuscript.

## Competing Interest

The authors declare no competing financial interest.

## References

1. Nadell CD, Xavier JB, Foster KR. 2008. The sociobiology of biofilms. FEMS Microbiology Reviews 33:206–224.

2. Allegrucci M, Hu FZ, Shen K, Hayes J, Ehrlich GD, Post JC, Sauer K. 2006. Phenotypic characterization of Streptococcus pneumoniaebiofilm development. J Bacteriol 188.

3. Ehrlich GD, Ahmed A, Earl J, Hiller NL, Costerton JW, Stoodley P, Post JC, DeMeo P, Hu FZ. 2010. The distributed genome hypothesis as a rubric for understanding evolution in situ during chronic bacterial biofilm infectious processes. FEMS Immunol Med Microbiol 59:269–79.

4. Ehrlich GD, Hiller NL, Hu FZ. 2008. What makes pathogens pathogenic. Genome biology 9:225.

5. Ehrlich GD, Hu FZ, Shen K, Stoodley P, Post JC. 2005. Bacterial plurality as a general mechanism driving persistence in chronic infections. Clin Orthop Relat Res:20–4.

6. Ito A, Taniuchi A, May T, Kawata K, Okabe S. 2009. Increased Antibiotic Resistance of Escherichia coli in Mature Biofilms. Applied and Environmental Microbiology 75:4093–4100.

7. Costerton W, Veeh R, Shirtliff M, Pasmore M, Post C, Ehrlich G. 2003. The application of biofilm science to the study and control of chronic bacterial infections. The Journal of clinical investigation 112:1466–1477.

8. Hall-Stoodley L, Nistico L, Sambanthamoorthy K, Dice B, Nguyen D, Mershon WJ, Johnson C, Hu FZ, Stoodley P, Ehrlich GD. 2008. Characterization of biofilm matrix, degradation by DNase treatment and evidence of capsule downregulation in Streptococcus pneumoniae clinical isolates. BMC microbiology 8:173.

9. Janto B, Ahmed A, Ito M, Liu J, Hicks DB, Pagni S, Fackelmayer OJ, Smith T-A, Earl J, Elbourne LDH, Hassan K, Paulsen IT, Kolstø A-B, Tourasse NJ, Ehrlich GD, Boissy R, Ivey DM, Li G, Xue Y, Ma Y, Hu FZ, Krulwich TA. 2011. Genome of alkaliphilic Bacillus pseudofirmus OF4 reveals adaptations that support the ability to grow in an external pH range from 7.5 to 11.4. Environmental Microbiology 13:3289–3309.

10. Braxton EE, Jr., Ehrlich GD, Hall-Stoodley L, Stoodley P, Veeh R, Fux C, Hu FZ, Quigley M, Post JC. 2005. Role of biofilms in neurosurgical device-related infections. Neurosurg Rev 28:249–55.

11. Wolcott RD, Ehrlich GD. 2008. Biofilms and Chronic Infections. JAMA 299:2682–2684.

12. Nguyen D, Joshi-Datar A, Lepine F, Bauerle E, Olakanmi O, Beer K, McKay G, Siehnel R, Schafhauser J, Wang Y, Britigan BE, Singh PK. 2011. Active starvation responses mediate antibiotic tolerance in biofilms and nutrient-limited bacteria. Science (New York, NY) 334:982–986.

13. Cashel M. 1969. The Control of Ribonucleic Acid Synthesis in Escherichia coli IV. RELEVANCE OF UNUSUAL PHOSPHORYLATED COMPOUNDS FROM AMINO ACID-STARVED STRINGENT STRAINS. The Journal of Biological Chemistry 244:3133–3141.

14. Haseltine WA, Block R. 1973. Synthesis of guanosine tetra- and pentaphosphate requires the presence of a codon-specific, uncharged transfer ribonucleic acid in the acceptor site of ribosomes. Proceedings of the National Academy of Sciences of the United States of America 70:1564–1568.

15. Cashel M. 1975. Regulation of bacterial ppGpp and pppGpp. Annual review of microbiology 29:301–318.

16. Nierlich DP. 1974. Regulation of bacterial growth. Science 184:1043–1050.

17. Erlich H, Laffler T, Gallant J. 1971. ppGpp formation in Escherichia coli treated with rifampicin. Journal of Biological Chemistry 246:6121–6123.

18. Agirrezabala X, Fernández IS, Kelley AC, Cartón DG, Ramakrishnan V, Valle M. 2013. The ribosome triggers the stringent response by RelA via a highly distorted tRNA. EMBO Reports 14:811–816.

19. Metzger S, Dror IB, Aizenman E, Schreiber G, Toone M, Friesen JD, Cashel M, Glaser G. 1988. The nucleotide sequence and characterization of the relA gene of Escherichia coli. J Biol Chem 263:15699–704.

20. Loveland AB, Bah E, Madireddy R, Zhang Y, Brilot AF, Grigorieff N, Korostelev AA. 2016. Ribosome•RelA structures reveal the mechanism of stringent response activation. eLife 5:e17029.

21. Arenz S, Abdelshahid M, Sohmen D, Payoe R, Starosta AL, Berninghausen O, Hauryliuk V, Beckmann R, Wilson DN. 2016. The stringent factor RelA adopts an open conformation on the ribosome to stimulate ppGpp synthesis. Nucleic Acids Res 44:6471–81.

22. Gratani FL, Horvatek P, Geiger T, Borisova M, Mayer C, Grin I, Wagner S, Steinchen W, Bange G, Velic A. 2018. Regulation of the opposing (p) ppGpp synthetase and hydrolase activities in a bifunctional RelA/SpoT homologue from Staphylococcus aureus. PLoS genetics 14:e1007514.

23. Ronneau S, Hallez R. 2019. Make and break the alarmone: regulation of (p) ppGpp synthetase/hydrolase enzymes in bacteria. FEMS microbiology reviews.

24. Hauryliuk V, Atkinson GC, Murakami KS, Tenson T, Gerdes K. 2015. Recent functional insights into the role of (p)ppGpp in bacterial physiology. Nature reviews Microbiology 13:298–309.

25. Wexselblatt E, Oppenheimer-Shaanan Y, Kaspy I, London N, Schueler-Furman O, Yavin E, Glaser G, Katzhendler J, Ben-Yehuda S. 2012. Relacin, a Novel Antibacterial Agent Targeting the Stringent Response. PLOS Pathogens 8:e1002925.

26. Wexselblatt E, Kaspy I, Glaser G, Katzhendler J, Yavin E. 2013. Design, synthesis and structure–activity relationship of novel Relacin analogs as inhibitors of Rel proteins. European Journal of Medicinal Chemistry 70:497–504.

27. Yanling C, Hongyan L, Xi W, Wim C, Dongmei D. 2018. Efficacy of relacin combined with sodium hypochlorite against Enterococcus faecalis biofilms. Journal of applied oral science : revista FOB 26:e20160608–e20160608.

28. Syal K, Flentie K, Bhardwaj N, Maiti K, Jayaraman N, Stallings CL, Chatterji D. 2017. Synthetic (p)ppGpp Analogue Is an Inhibitor of Stringent Response in Mycobacteria. Antimicrobial Agents and Chemotherapy 61:e00443–17.

29. Dutta NK, Klinkenberg LG, Vazquez M-J, Segura-Carro D, Colmenarejo G, Ramon F, Rodriguez-Miquel B, Mata-Cantero L, Porras-De Francisco E, Chuang Y-M, Rubin H, Lee JJ, Eoh H, Bader JS, Perez-Herran E, Mendoza-Losana A, Karakousis PC. 2019. Inhibiting the stringent response blocks <em>Mycobacterium tuberculosis</em> entry into quiescence and reduces persistence. Science Advances 5:eaav2104.

30. de la Fuente-Núñez C, Reffuveille F, Haney EF, Straus SK, Hancock REW. 2014. Broad-Spectrum Anti-biofilm Peptide That Targets a Cellular Stress Response. PLoS Pathog 10:e1004152.

31. Pletzer D, Hancock RE. 2016. Antibiofilm Peptides: Potential as Broad-Spectrum Agents. J Bacteriol 198:2572–8.

32. Dostert M, Belanger CR, Hancock REW. 2019. Design and Assessment of Anti-Biofilm Peptides: Steps Toward Clinical Application. Journal of Innate Immunity 11:193–204.

33. Mansour SC, de la Fuente-Núñez C, Hancock REW. 2015. Peptide IDR-1018: modulating the immune system and targeting bacterial biofilms to treat antibiotic-resistant bacterial infections. Journal of Peptide Science 21:323–329.

34. Andresen L, Tenson T, Hauryliuk V. 2016. Cationic bactericidal peptide 1018 does not specifically target the stringent response alarmone (p)ppGpp. Scientific Reports 6:36549.

35. Wadood A, Ahmed N, Shah L, Ahmad A, Hassan H, Shams S. 2013. In-silico drug design: An approach which revolutionarised the drug discovery process. OA drug design & delivery 1:3–7.

36. Brown A, Fernández IS, Gordiyenko Y, Ramakrishnan V. 2016. Ribosome-dependent activation of stringent control. Nature 534:277–280.

37. Manav MC, Beljantseva J, Bojer MS, Tenson T, Ingmer H, Hauryliuk V, Brodersen DE. 2018. Structural basis for (p)ppGpp synthesis by the Staphylococcus aureus small alarmone synthetase RelP. J Biol Chem 293:3254–3264.

38. Steinchen W, Vogt MS, Altegoer F, Giammarinaro PI, Horvatek P, Wolz C, Bange G. 2018. Structural and mechanistic divergence of the small (p)ppGpp synthetases RelP and RelQ. Sci Rep 8:2195.

39. Hogg T, Mechold U, Malke H, Cashel M, Hilgenfeld R. 2004. Conformational Antagonism between Opposing Active Sites in a Bifunctional RelA/SpoT Homolog Modulates (p)ppGpp Metabolism during the Stringent Response. Cell 117:57–68.

40. Singal B, Balakrishna AM, Nartey W, Manimekalai MSS, Jeyakanthan J, Gruber G. 2017. Crystallographic and solution structure of the N-terminal domain of the Rel protein from Mycobacterium tuberculosis. FEBS Lett 591:2323–2337.

41. Lipinski CA, Lombardo F, Dominy BW, Feeney PJ. 1997. Experimental and computational approaches to estimate solubility and permeability in drug discovery and development settings. Advanced Drug Delivery Reviews 23:3–25.

42. Sliwoski G, Kothiwale S, Meiler J, Lowe EW. 2014. Computational methods in drug discovery. Pharmacological reviews 66:334–395.

43. Król JE, Hall DC, Balashov S, Pastor S, Sibert J, McCaffrey J, Lang S, Ehrlich RL, Earl J, Mell JC, Xiao M, Ehrlich GD. 2019. Genome rearrangements induce biofilm formation in Escherichia coli C – an old model organism with a new application in biofilm research. BMC Genomics 20:767.

44. Payoe R, Fahlman RP. 2011. Dependence of RelA-Mediated (p)ppGpp Formation on tRNA Identity. Biochemistry 50:3075–3083.

45. Åberg A, Fernández-Vázquez J, Cabrer-Panes JD, Sánchez A, Balsalobre C. 2009. Similar and divergent effects of ppGpp and DksA deficiencies on transcription in Escherichia coli. Journal of bacteriology 191:3226–3236.

46. Blattner FR, Plunkett G, Bloch CA, Perna NT, Burland V, Riley M, Collado-Vides J, Glasner JD, Rode CK, Mayhew GF. 1997. The complete genome sequence of Escherichia coli K-12. science 277:1453–1462.

47. Olins PO, Nomura M. 1981. Regulation of the S10 ribosomal protein operon in E. coli: nucleotide sequence at the start of the operon. Cell 26:205–211.

48. Lemke JJ, Sanchez-Vazquez P, Burgos HL, Hedberg G, Ross W, Gourse RL. 2011. Direct regulation of Escherichia coli ribosomal protein promoters by the transcription factors ppGpp and DksA. Proceedings of the National Academy of Sciences 108:5712–5717.

49. Burgos HL, O’Connor K, Sanchez-Vazquez P, Gourse RL. 2017. Roles of transcriptional and translational control mechanisms in regulation of ribosomal protein synthesis in Escherichia coli. Journal of bacteriology 199:e00407–17.

50. Gawin A, Peebo K, Hans S, Ertesvåg H, Irla M, Neubauer P, Brautaset T. 2019. Construction and characterization of broad-host-range reporter plasmid suitable for on-line analysis of bacterial host responses related to recombinant protein production. Microbial cell factories 18:80.

51. Mechold U, Cashel M, Steiner K, Gentry D, Malke H. 1996. Functional analysis of a relA/spoT gene homolog from Streptococcus equisimilis. Journal of Bacteriology 178:1401–1411.

52. Wendrich TM, Blaha G, Wilson DN, Marahiel MA, Nierhaus KH. 2002. Dissection of the mechanism for the stringent factor RelA. Molecular cell 10:779–788.

53. Shankarappa B, Sirko D, Ehrlich G. 1992. A general method for the identification of regions suitable for site-directed silent mutagenesis. Biotechniques 12:382–384.

54. Shankarappa B, Vijayananda K, Ehrlich G. 1992. SILMUT: a computer program for the identification of regions suitable for silent mutagenesis to introduce restriction enzyme recognition sequences. Biotechniques 12:882–884.

55. Harder E, Damm W, Maple J, Wu C, Reboul M, Xiang JY, Wang L, Lupyan D, Dahlgren MK, Knight JL. 2016. OPLS3: a force field providing broad coverage of drug-like small molecules and proteins. Journal of chemical theory and computation 12:281–296.

56. Halgren TA, Murphy RB, Friesner RA, Beard HS, Frye LL, Pollard WT, Banks JL. 2004. Glide: A New Approach for Rapid, Accurate Docking and Scoring. 2. Enrichment Factors in Database Screening. Journal of Medicinal Chemistry 47:1750–1759.

57. Friesner RA, Banks JL, Murphy RB, Halgren TA, Klicic JJ, Mainz DT, Repasky MP, Knoll EH, Shelley M, Perry JK, Shaw DE, Francis P, Shenkin PS. 2004. Glide: A New Approach for Rapid, Accurate Docking and Scoring. 1. Method and Assessment of Docking Accuracy. Journal of Medicinal Chemistry 47:1739–1749.

58. Lipinski CA, Lombardo F, Dominy BW, Feeney PJ. 2001. Experimental and computational approaches to estimate solubility and permeability in drug discovery and development settings1. Advanced Drug Delivery Reviews 46:3–26.

59. Lipinski CA. 2004. Lead-and drug-like compounds: the rule-of-five revolution. Drug Discovery Today: Technologies 1:337–341.

60. Benet LZ, Hosey CM, Ursu O, Oprea TI. 2016. BDDCS, the Rule of 5 and drugability. Advanced Drug Delivery Reviews 101:89–98.

61. Bickerton GR, Paolini GV, Besnard J, Muresan S, Hopkins AL. 2012. Quantifying the chemical beauty of drugs. Nat Chem 4:90–98.

62. Hall DC, Ji H-F. 2020. A search for medications to treat COVID-19 via in silico molecular docking models of the SARS-CoV-2 spike glycoprotein and 3CL protease. Travel Medicine and Infectious Disease doi:https://doi.org/10.1016/j.tmaid.2020.101646:101646.

63. Potrykus K, Murphy H, Philippe N, Cashel M. 2011. ppGpp is the major source of growth rate control in E. coli. Environmental microbiology 13:563–575.

64. Gaal T, Gourse RL. 1990. Guanosine 3’-diphosphate 5’-diphosphate is not required for growth rate-dependent control of rRNA synthesis in Escherichia coli. Proceedings of the National Academy of Sciences 87:5533–5537.

65. Hernandez VJ, Bremer H. 1993. Characterization of RNA and DNA synthesis in Escherichia coli strains devoid of ppGpp. Journal of Biological Chemistry 268:10851–10862.

66. Stoitsova SR, Paunova-Krasteva TS, Borisova DB. 2016. Modulation of biofilm growth by sub-inhibitory amounts of antibacterial substances. Microbial Biofilms-Importance and Applications.

67. Jaroslaw Krol DH, Garth Ehrlich. 2018. ASM Biofilms JEKmagTech 2018 abstr ASM Biofilms 2018 Washington DC •

68. Ji H-F, Ehrlich GD, Hall Jr DC, Krol JE, Cahill JP. 2020. RelA Inhibitors for Biofilm Disruption. Google Patents.

69. Madhavi Sastry G, Adzhigirey M, Day T, Annabhimoju R, Sherman W. 2013. Protein and ligand preparation: parameters, protocols, and influence on virtual screening enrichments. Journal of Computer-Aided Molecular Design 27:221–234.

70. Shankarappa B, Balachandran R, Gupta P, Ehrlich G. 1992. Introduction of multiple restriction enzyme sites by in vitro mutagenesis using the polymerase chain reaction. Genome Research 1:277–278.

71. Kitagawa M, Ara T, Arifuzzaman M, Ioka-Nakamichi T, Inamoto E, Toyonaga H, Mori H. 2005. Complete set of ORF clones of Escherichia coli ASKA library (A Complete S et of E. coli K-12 ORF A rchive): Unique Resources for Biological Research. DNA research 12:291–299.

72. Krol JE, Biswas S, King C, Biswas I. 2014. SMU.746-SMU.747, a putative membrane permease complex, is involved in aciduricity, acidogenesis, and biofilm formation in Streptococcus mutans. J Bacteriol 196:129–39.

